# The reindeer circadian clock is rhythmic and temperature-compensated but shows evidence of weak coupling between the secondary and core molecular clock loops

**DOI:** 10.1101/2024.04.24.590752

**Authors:** Daniel Appenroth, Chandra S Ravuri, Sara K Torppa, Shona H Wood, David G Hazlerigg, Alexander C West

**Affiliations:** Arctic Seasonal Timekeeping Initiative (ASTI), Arctic Chronobiology and Physiology research group, Department of Arctic and Marine Biology, UiT — The Arctic University of Norway, Framstredet 42, 9019 Tromsø, Norway

**Keywords:** circadian, polar, reindeer, *Rangifer tarandus*, temperature-compensation, entrainment, cell culture

## Abstract

Circadian rhythms synchronize the internal physiology of animals allowing them to anticipate daily changes in their environment. Arctic habitats may diminish the selective advantages of circadian rhythmicity by relaxing daily rhythmic environmental constraints, presenting a valuable opportunity to study the evolution of circadian rhythms. In reindeer, circadian control of locomotor activity and melatonin release is weak or absent, and the molecular clockwork is reportedly non-functional. Here we present new evidence that the circadian clock in cultured reindeer fibroblasts is rhythmic and temperature-compensated. Compared to mouse fibroblasts, however, reindeer fibroblasts have a short free-running period, and temperature cycles have an atypical impact on clock gene regulation. In reindeer cells, *Per2* and *Bmal1* reporters show rapid responses to temperature cycles, with a disintegration of their normal antiphasic relationship. The antiphasic *Per2* - *Bmal1* relationship re-emerges immediately after release from temperature cycles, but without complete temperature entrainment and with a marked decline in circadian amplitude. Experiments using *Bmal1* promoter reporters with mutated RORE sites showed that a reindeer-like response to temperature cycles can be mimicked in mouse or human cell lines by decoupling *Bmal1* reporter activity from ROR / REV-ERB dependent transcriptional regulation. We suggest that weak coupling between core and secondary circadian feedback loops accounts for the observed behaviour of reindeer fibroblasts *in vitro*. Our findings highlight diversity in how the thermal environment affects the temporal organisation of mammals living under different thermoenergetic constraints.

## INTRODUCTION

In mammals, circadian rhythms in behaviour and physiology are coordinated by a cell-autonomous molecular clock (Takahashi, 2017). The principal mechanism of the molecular clock begins with the activation of transcription by heterodimers formed of the clock genes *Clock* and *Bmal1*. The CLOCK:BMAL1 heterodimer initiates transcription by binding to E-box elements, including those found in the promotors of the clock gene families of *Per* and *Cry*. Once translated, PER and CRY proteins heterodimerise and translocate to the nucleus where they block the transactivation potential of CLOCK:BMAL1, bringing a halt to their own transcription and ultimately resetting the circadian cycle (Takahashi, 2017). This core loop is stabilised by a secondary loop in which retinoic acid-related orphan receptors (RORs) and REV-ERBs respectively stimulate and inhibit *Bmal1* transcription *via* ROR response elements (ROREs) (Abe et al., 2022; Akashi and Takumi, 2005; Guillaumond et al., 2005; Preitner et al., 2002; Ueda et al., 2002). The expression of the *Ror* and *Rev-erb* genes are, in part, under the control of CLOCK:BMAL1 which regulates expression through E-box elements (Preitner et al., 2002) thereby coupling the core and secondary loops together, maintaining robust circadian oscillation.

The adaptive value of circadian rhythms is thought to be linked to the synchronisation between the organism and rhythmic changes in the environment associated with day and night (West and Bechtold, 2015). This leads to questions regarding the role of circadian rhythms in weakly rhythmic environments. The polar regions are defined by the reduced amplitude of the daily solar cycle, leading to continuous periods of darkness in the winter (polar night) and of continuous daylight during the summer (polar day). Under such conditions, daily rhythms in environmental factors are of reduced amplitude, and strictly enforced daily partitioning of behaviour and physiology through the circadian system might restrain the organism’s ability to exploit favourable conditions (Hazlerigg et al., 2023).

The reindeer (*Rangifer tarandus*) has received significant attention as a model for polar circadian biology (Arnold et al., 2018; Erriksson et al., 1981; Lin et al., 2019; Loe et al., 2007; Lu et al., 2010; Meier et al., 2024; Stokkan et al., 2007; van Oort et al., 2007). Throughout the year, reindeer activity patterns show a strong ultradian component, reflecting ruminant feeding behaviour, and during the polar night and polar day, the daily (24-h) component of their activity patterns diminishes to low or undetectable levels (Arnold et al., 2018; van Oort et al., 2005, 2007). Plasma concentration of melatonin, a hormone whose release is under circadian control in most mammals, is acutely dark-inducible in reindeer and shows no evidence of circadian rhythms under constant conditions (Stokkan et al., 2007). Most intriguingly, an *in vitro* study of the reindeer circadian clock reported that cultured reindeer skin fibroblasts transduced with *Bmal1:luc* and *Per2:luc* reporter constructs had weak, rapidly dampening rhythms with large variation in period (19-31h) (Lu et al., 2010). More recently, a reindeer-specific point mutation in the *Per2* gene which reduces the heterodimerisation potential of reindeer-like PER2 with CRY1, has been suggested as an explanation for the reported weak molecular clock of reindeer (Lin et al., 2019).

Our objective at the start of this study was to understand the mechanistic basis of the weak circadian rhythmicity of reindeer skin fibroblasts. For this purpose, we first attempted to replicate the results reported by Lu et al. (2010). We established primary cultures of mouse and reindeer skin fibroblasts and transduced them with either *Bmal1:luc* or *Per2:luc* promoter reporters. Against our expectations, we found temperature-compensated, rhythmic reporter activity in reindeer fibroblasts. Follow-up experiments revealed an atypical response to a temperature entrainment paradigm in reindeer fibroblasts, with acute sensitivity of both *Bmal1* and *Per2* to temperature increments apparently working against efficient entrainment. We suggest that these results may be due to a weakened coupling between the core and secondary loops of the circadian clock machinery.

## MATERIAL AND METHODS

### Animals

All animals were kept in accordance with the EU directive 201/63/EU under licenses provided by the Norwegian food safety authority (Mattilsynet, FOTS 28929). Reindeer-derived biological material stemmed exclusively from a semi-domesticated herd held at the University of Tromsø, Norway (69°N).

### Fibroblast cell cultures

Fibroblasts were cultivated from an ear skin sample of euthanized reindeer (*Rangifer tarandus tarandus*, female, 9 months old) and from an ear puncture of a laboratory mouse (*Mus musculus*, C57Bl6/J, males, 10 weeks old, provided by Simón Santamaría and Karolina Szafranska, UiT). Sampling and cultivation of fibroblasts broadly followed the protocols outlined in Du and Brown (2021).

In brief, tissue samples were collected in sampling medium containing Dulbecco’s Modified Eagle Medium (DMEM, Merck D5796), 50% fetal bovine serum (FBS, Biowest S181B), 1% penicillin-streptomycin (P/S, Merck P4333) and 1x amphotericin B on ice, then incubated in 2ml digestion medium containing DMEM, 10% FBS, 1% P/S, 1x amphoterecin B and 3.5U liberase for 4h at 37°C and 5% CO_2_. Digested fragments were transferred into a 50 ml falcon tube filled with warm phosphate-buffered saline (PBS, Biowest L0615) and centrifuged for 5 min at 200 x g. The supernatant was removed and the formed pellet was resuspended in 0.2 ml culture medium (DMEM + 10% FBS + 1% P/S). The mix was placed in the centre of a fresh 6-well plate, overlayed with a Millicell cell culture insert (Merck, PICM0RG50) and provided with 2 ml culture medium. The sample was incubated for ca. 2 weeks at 37°C and 5% CO_2_ with a medium change every 3-4 days. Ready-to-harvest cells were trypsinized (Merck T4049) and replated. Cells were passaged until an appropriate volume was achieved. Active cultures were kept in DMEM, 10% FBS and 1% P/S and housed in cell incubators at 37°C, 5% CO_2_ and high humidity. Cultures not in active use were stored in liquid nitrogen in culture medium + 10% dimethyl sulfoxide (DMSO, Merck C6164).

When required, frozen cells were thawed quickly by incubating them in a water bath at 37°C. Thawed cells were mixed with 10ml culture medium, mixed and centrifuged for 5 minutes at 300 x g. The supernatant was discarded and the pellet was resuspended in 7 ml culture medium. Resuspended cells were transferred into a cell culture flask and incubated at 37°C and 5% CO_2._

### Generation of the *Per2:luc* reporter

Circadian clock gene activity was recorded with a *Bmal1:luc* reporter (Addgene 68833, a gift from Steven Brown) and a *Per2:luc* reporter (Addgene 212035) contained in lentiviral transfer vectors (referred to as pLV6-Bmal-luc vector and pLV6-Per2-luc vector respectively). The *Bmal1:luc* reporter contains a murine *Bmal1* promoter with an adjacent *luciferase* sequence. The *Per2:luc* reporter was constructed in-house using a restriction enzyme strategy, which is efficient in constructing large plasmids (Zhang and Tandon, 2012). The *Per2:luc* reporter vector (pLV6-Per2-luc) consists of the backbone of the *Bmal1:luc* reporter vector (pLV6-Bmal-luc) and a murine *Per2* promoter with adjacent *luciferase* sequence which is contained in the pGL3 basic E2 vector (Addgene 48747, a gift from Joseph Takahashi). The QuickChange Lightning Site-Directed Mutagenesis (SDM) kit (Agilant 210518) following the manufacturer’s manual was used to ligate the *Per2:luc* reporter with pLV6 backbone as follows.

The *Bmal1:luc* reporter vector contains one BamHI cutting site 3’ of the *Bmal1:luc* insert. The SDM kit and custom-designed oligonucleotides (**Suppl. Table S1**) were used to create an additional BamHI cutting site 5’ of the insert by a single nucleotide mutation. After the SDM kit procedure, transformation of the mutated plasmid was done with stable competent E. coli (NEB C3040H) according to the manufacturer’s manual. Cells were grown on LB agar plates (40g/l LB-agar, 100 µg/ml Ampicillin (Merck A9393)) and single clones were cultivated in LB broth overnight (25g LB, 100 µg/ml Ampicillin). DNA was extracted using the Qiagen miniprep (Qiagen 12123) and the mutation was checked by whole plasmid sequencing (Plasmidsaurus). After confirmation, the *Bmal1:luc* reporter vector was digested with BamHI (NEB R3136S), supplemented with calf intestinal alkaline phosphatase (Quick CIP, NEB M0525S) to avoid self-ligation. The digest was run on an 1% Agarose gel (containing ethidium bromide) and the pLV6 backbone was extracted with the QIAquick Gel extraction kit (Qiagen 28704) according to the manufacturer’s manual.

Conveniently, the murine *Per2* promoter and *luciferase* sequence in the pGL3-Per2-luc vector is by default flanked by BamHI cutting sites. Consequently, the pGL3-Per2-luc vector was directly digested with BamHI and the *Per2:luc* sequence was extracted from an agarose gel as explained above.

After successful digestions and extractions, the pLV6 backbone of the *Bmal1:luc* reporter vector and the *Per2:luc* fragment were ligated with T4 DNA ligase (NEB M0202S) according to the manufacturer’s manual. Subsequently, extracted vectors from transfected single colonies were sequenced by Plasmidsaurus and checked for correct ligation and correct sequence.

### Generation of the *Bmal1:luc* with mutated RORE sites

The two RORE elements (designated RORE1 and RORE2) of the *Bmal1:luc* reporter were mutated in series using the SDM kit following the manufacturer’s instructions. RORE-specific primers (**Suppl. Table S1**) were designed to mutate RORE sequences as previously reported (Guillaumond et al., 2005). Amplification of plasmids was achieved as described above. Confirmation of RORE mutation was confirmed by whole plasmid sequencing (Plasmidsaurus). The resulting reporter is subsequently referred to as *Bmal1-mutated:luc* and the whole vector is referred to as pLV6-Bmal1-mut-luc.

### Lentiviral vector production and transduction

The lentiviral vectors containing the respective reporters (*Bmal1:luc*, *Per2:luc* and *Bmal1-mutated:luc*) were packed following broadly Du and Brown (2021) and Salmon and Trono (2007).

The day before transfection, HEK293T/17 cells (ATCC CRL-11268) were seeded at a density of 2×10^6^ /10cm dish with the aim of a confluency of 40-50%. Packaging plasmids pMD2G and psPAX2 (Addgene 12259 and 12260, gifts from Didier Trono) and the respective transfer vector (pLV6-Bmal-luc, pLV6-Per2-luc or pLV6-Bmal-mut-luc) were dissolved in TE buffer, p.H 8.0 (Qiagen 12162) to a total volume of 250µl. Next, 500µl HEPES-buffered saline solution (Alfa Aesar AAJ62623AK) was added. 250 µl of 0.5M CaCl_2_ were added in an empty sterile 15 ml tube. The DNA/ TE/ HeB mix was added dropwise into the CaCl_2_ containing tube under constant vortexing. The mixture was then incubated for 20-30 minutes at room temperature during which a fine white translucent precipitate formed. 1 ml of the precipitate was dropwise added to the fibroblast culture and gently mixed. The next morning, the old medium was removed, washed twice with PBS and 10 ml of fresh culture medium was added. For the next two days 10 ml of culture medium was collected and replenished afterwards. The pooled collection medium from two consecutive days (in total 20 ml) was centrifuged for 5 min at 500x g and 4°C. Thereafter the supernatant was filtered through 0.45-um syringe filters and then stored as 1 ml aliquots at –80°C until further use.

Cultivated mouse and reindeer fibroblast were transduced with lentivirus containing either of the transfer vectors. The day before infection, confluent fibroblasts were split to achieve 50% confluency on the following day. The medium was replaced with warmed lentivirus-containing medium supplemented with 8 µg /ml protamine sulfate. After 48 h the lentivirus-containing medium was renewed and after three days the medium was replaced with a medium containing blasticidin selection antibiotic (Merck 15205). The cells were then grown in T75 culture flasks before seeding into 35mm dishes and were prepared for recording.

2-3 days before the recording the cultures were transferred into a recording medium containing 5% FBS and 0.1 mM luciferin (Promega E1601) (Feeney et al., 2016; Yamazaki and Takahashi, 2005). Synchronisation of the cultures was achieved by dexamethasone (DEX) (Merck D4902). Recording media in the culture dishes was exchanged for synchronization medium containing DEX (Recording media + 100nM DEX) and incubated for ca. 30 minutes in the culture incubator (37°C, 5% CO_2_). After incubation, the cultures were washed with PBS twice and a final volume of 2-3 ml recording media was filled in the 35 mm culture dishes. The dishes were sealed with parafilm and placed into a photomultiplier tube (PMT) (Hamamatsu Photonics LM-2400) placed inside a cell incubator. The PMT was connected to a PC with data acquisition software.

### Cell culture experiments (including data analysis)

#### Experiment 1: Different temperatures, luminescence recording

Mouse and reindeer fibroblast cultures were either transduced with the clock gene promoter-reporter *Bmal1:luc* (pLV6-Bmal1-luc) or *Per2:luc* (pLV6-Per2-luc). Fibroblast cultures were first measured under different temperatures in the following order 40 °C, 34 °C, 31 °C and 37 °C. Luminescence was measured for 4-6 days under each temperature. Between each temperature recording, all cultures were re-synchronized as outlined above. Six replicates were used for each species-reporter combination, accumulating to a total of 24 cultures. However, 2 replicates of the reindeer-*Per2* culture died early in the experiment resulting in 4 replicates for this group.

For each replicate, moving averages were calculated for 24 hours around each respective data point, i.e. 12 hours before and after. Phases were assessed by calculating the centre of gravity of the bioluminescent peak on the first day 24h after DEX synchronisation. The centre of gravity was generated by the CircWave programme (version 1.4, programmed by Dr. Roelof A. Hut). Phases were then normalized against the free-running period of the respective species and plotted in a circular graph in R studio (package: ggplot2). The displayed values represent means from all replicates recorded at 37°C.

Periods, half-life times and Q10 values were calculated based on damped sine wave analysis fitted with GraphPad prism (version 9.4.0) to *Bmal1:luc* recordings of mouse and reindeer cultures. Periods and half-life times were extracted from the damped sine wave analysis for each temperature. Periods under different temperatures were used to calculate the Q10 value in R studio (R package: respirometry). Figures were produced in GraphPad prism. Statistical tests were performed in GraphPad prism.

#### Experiment 2: Temperature entrainment, luminescence recording

In the second experiment, the ability of temperature entrainment for each species and each gene was measured. For each group, three replicates were used and transferred into the PMT under constant 37.2°C. Approximately 12 hours after the first synchronization, another three replicates of each group were synchronized. This resulted in half of each group being in anti-phase to the other half. The cultures were recorded for two days under constant 37.2°C. Thereafter the cultures were transferred into a temperature cycle for four days with 12h under 40.2 °C and 12h under 37.2 °C. After those four days, the cultures were again transferred to 37.2°C and measured for 2 days. Temperature was simultaneously recorded by a temperature logger (iButton, Maxim Integrated DS2922L) placed in the PMT and temperature was extracted with the programme OneWireViewer (Maxim Integrated Version 0.3.19.47).

Moving averages were calculated as described above. Centres of gravity were determined for each day of the recording in mouse and reindeer fibroblast and plotted as Zeitgeber time (ZT = 0 refers to the first uprise in temperature). The values of the last day of the recording were compared by an unpaired t-test. All graphs were produced in GraphPad prism.

#### Experiment 3: Serial collection of mouse and reindeer skin fibroblasts for qPCR analysis

Mouse and reindeer skin fibroblasts were seeded into 6 well plates and grown to confluency at 37.2°C in DMEM + 10% FBS + P/S. Cultures were then synchronised using 100nM DEX for 30 minutes and returned to the incubator for 48h. After 50h, we increased the temperature of the incubator to 40.2°C for 12h before returning the temperature to 37°C until the end of the experiment. Plates were sampled starting at 48h after synchronisation and then every four hours for the next 24h. During sampling, the plates were rapidly retrieved from the incubator, the media was then removed before the dishes were washed 3x in ice-cold PBS. Finally, all excess PBS was removed before the plates were snap-frozen on dry ice and transferred to –80°C freezer for storage.

RNA was extracted from each plate using RNeasy mini kit (QIAgen) and QIAshredder following the manufacturer’s instructions. We next converted 2 ug of RNA to cDNA using a high-capacity RNA to cDNA kit (Thermofisher) and performed qPCR using GoTaq qPCR reagent. Gene and species-specific qPCR primers (**Suppl. Table S1**) were designed from genomic sequences (mouse, GRCm39; reindeer, GCA_949782905.1). We confirmed primer specificity by cloning and sequencing of PCR product and ensured primer efficiency (90-105%) by standard dilutions. The housekeeping gene *Ppib* was used for data normalisation.

#### Experiment 4: Temperature entrainment of RORE mutated Bmal1 reporter

In this experiment, U2OS cells, mouse fibroblasts and reindeer fibroblasts were either transduced with the normal *Bmal1:luc* (pLV6-Bmal1-luc) or the *Bmal1-mutated:luc* (pLV6-Bmal1-mut-luc). First, those cultures were recorded under constant conditions for 2 days. Thereafter, a three-stage protocol of temperature entrainment similar to experiment 2 was applied, with recordings under constant ambient temperature, followed by 4 days of a temperature cycle, followed by a release back into constant temperature. Moving averages were calculated as described above. Graphs were plotted in GraphPad.

### Comparison of reindeer and mouse *Bmal1* promoter sequences

Genomic comparison was conducted by aligning the *Bmal1* promoter sequence of the *Bmal1:luc* reporter (pLV6-Bmal1-luc) to the *Bmal1* promoter sequence in reindeer (GCA_004026565.1) with Clustal Omega (https://www.ebi.ac.uk/). Furthermore, amino acid sequences of RORs and REV-ERBs between mouse and reindeer were compared by aligning them with Clustal Omega.

## RESULTS

### Clock gene promoter reporters are rhythmic and temperature-compensated in reindeer skin fibroblasts

We established primary cultures of mouse and reindeer skin fibroblasts and transduced them with either *Bmal1:luc* or *Per2:luc* clock gene promoter reporters. We then synchronised the cells with dexamethasone (DEX) and monitored bioluminescence of the reporter constructs for several days. In contrast to earlier work (Lu et al., 2010), both mouse and reindeer skin fibroblasts showed strong evidence of circadian oscillations, as shown by the reporter activity of *Bmal1:luc* and *Per2:luc* (**Figure 1a**). The free running period (τ) of the *Bmal1:luc* was 25h 20min in mouse and 21h 19min in reindeer. Both species, however, had near-identical phase angles (Ψ) between the *Per2* and *Bmal1* reporters (**Figure 1a, right panels**).

**Figure 1.**
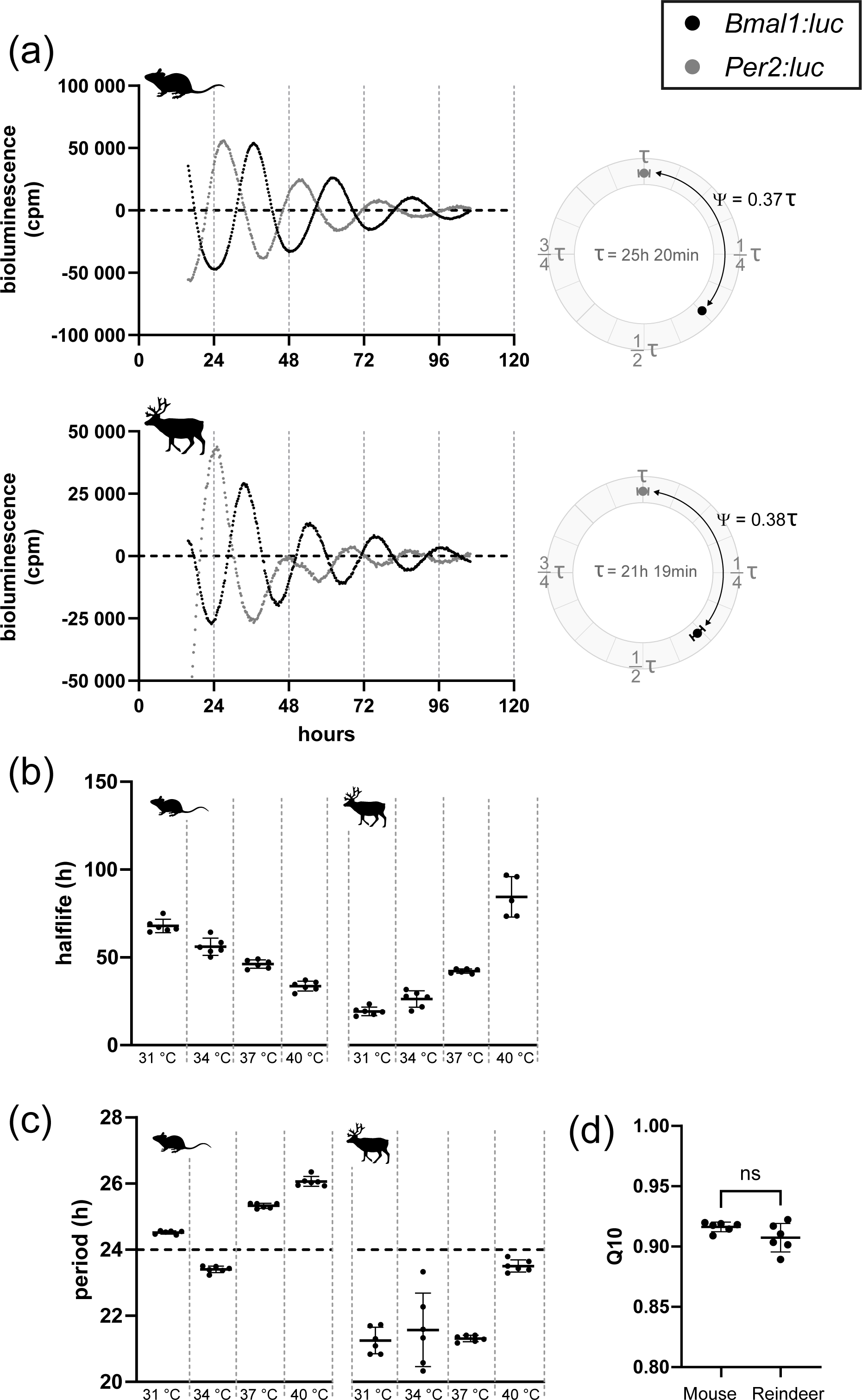
Endogenous and temperature-compensated clock gene expression in mouse and reindeer fibroblast culture. (a) Representative luminescence data of mouse and reindeer fibroblasts under 37°C ambient temperature. Expression of the *Bmal1*:luc reporter (black) is in approximate antiphase to the expression *of Per2:luc* reporter (grey) in both mouse and reindeer fibroblast cultures. Bioluminescence was measured in counts per minute (cpm). Phase angle difference (Ψ) between *Bmal1*:luc and *Per2:luc* for the first day after DEX synchronisation (from hour 24 to hour 48) are plotted as circular plots. Data is normalized for the respective free-running period (τ). (b) Half-life times of *Bmal1*:luc expression under different ambient temperatures. Half-life times were calculated by damped sine wave analysis. (c) Periods of *Bmal1*:luc expression under different ambient temperatures. Periods have been determined by damped sine wave analysis. (d) Q10 values of *Bmal1*:luc in mouse and reindeer fibroblast cultures. Q10 values are based on periods as shown in panel c. Both Q10 values are within the acceptable range for temperature-compensation (0.8-1.4) and do not differ significantly from each other (ns).

Besides sustained rhythmicity under constant conditions, circadian oscillators are defined by their stable period length at different temperatures, a quality known as temperature compensation (Sweeney and Hastings, 1960). To test temperature-compensation, and hence the robustness, of mouse and reindeer skin fibroblasts we measured the oscillatory periods of cells transduced with the *Bmal1:luc* promoter reporter in cells incubated at 31°C, 34°C, 37°C or 40°C (**Suppl. Fig. S1**). Although incubation temperature changed the dampening half-lives of the *luciferase* reporter (One-way ANOVA, F_mouse_= 97.73, F_reindeer_ = 126.9, p for both < 0.0001) (**Figure 1b**), both mouse and reindeer skin fibroblast cells remained rhythmic at all temperatures and maintained stable period lengths (**Figure 1c**). Q10 values were similar in mouse (0.916) and reindeer cells (0.907) (t-test, p = 0.108, t = 1.766, df = 10), and well within the definitive 0.8-1.4 range of temperature-compensation (**Figure 1d**) (Sweeney and Hastings, 1960).

### Reindeer cells show weak entrainment under 24h temperature cycles

Daily fluctuations in body temperature are thought to be a universal entrainment cue in mammals (Buhr et al., 2010). To test temperature synchronisation in mouse and reindeer skin fibroblasts we ran a three-stage experiment. In stage 1, mouse and reindeer skin fibroblasts, transduced with either *Bmal1:luc* or *Per2:luc* reporters, were DEX-synchronised at two time points 12h apart. Bioluminescence of those cultures was recorded under a constant temperature until stage 2. In stage 2, we exposed the cultures to a fluctuating temperature cycle with an amplitude of 3°C for four 24h cycles. In stage 3, the cultures were released back to constant temperature. The prediction for a temperature-entrainable circadian clock is that in stage 1 the cultures show two distinct phases 12h apart, stage 2 will entrain cultures into the same phase, and stage 3 will reveal that this new phase persists in constant conditions.

In mouse fibroblasts, the *Per2:luc* reporter (**Figure 2a**) responded immediately to the temperature cycles (**Figure 2c and Suppl. Fig. S2b**) and the phase of the 12h-apart synchronized cultures remained in phase to each other once back on constant temperature (t-test, p = 0.0856, t = 2.271, df = 4). For the *Bmal1:luc* reporter (**Figure 2b**), the temperature cycle revealed a gradual entrainment (**Figure 2c and Suppl. Fig. S2b**) which emerged over several days. Release of the cultures from a temperature cycle to constant temperature showed near-synchrony of cultures whose acrophases (i.e. centre of gravity) were 44 min ± 13 min apart from each other (**Figure 2c**) (t-test, p = 0.0288, t = 3.341, df = 4). These data are consistent with previous temperature entrainment experiments in mouse cells (Saini et al., 2012).

**Figure 2.**
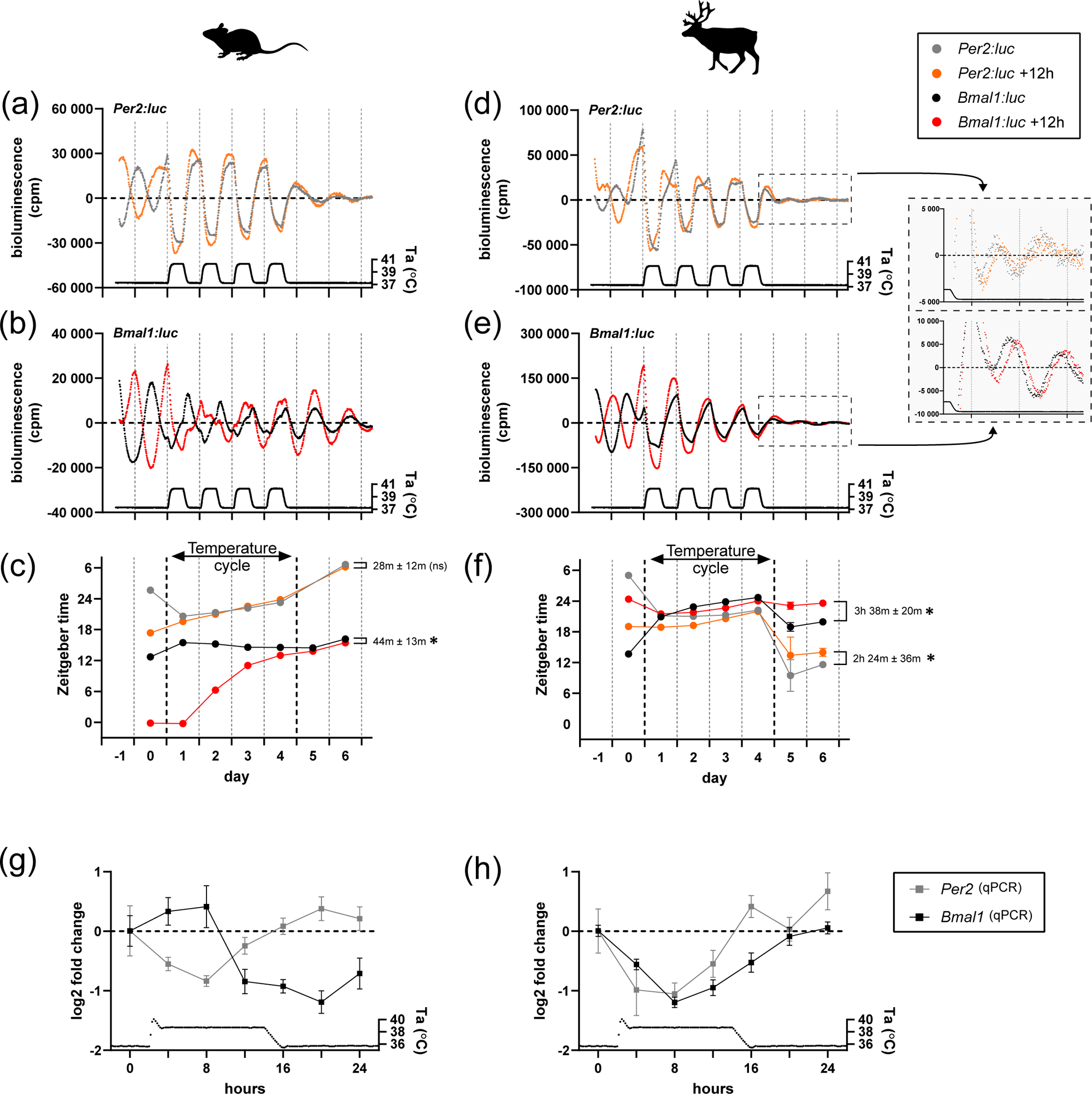
Temperature entrainment of clock genes expression in mouse and reindeer fibroblast cultures. Expression of *Per2:luc* and *Bmal1:luc* in mouse (a-c) and reindeer fibroblasts (d-f) under constant and cycling ambient temperature (Ta). Two sets of cultures were DEX-synchronized 12h apart from each other. Bioluminescence was measured in counts per minute (cpm). For reindeer, bioluminescence data after the temperature cycle were plotted in zoomed-in plots beside the corresponding graphs. Times of peak expressions (centre of gravity) were determined with the CircWave programme for each rhythm to track phase relationships and responses before, during and after the temperature cycle (c, f). Phase relationships between the initially 12h-apart synchronised cell cultures of the same gene were calculated after the temperature cycle and tested with an unpaired t-test (asterisk indicates p< 0.05, ns denotes p> 0.05). (g, h) Relative gene expressions of *Bmal1* and *Per2* in mouse and reindeer fibroblasts under one temperature cycle. Gene expression data was measured by qPCR with species-specific primers.

In reindeer fibroblasts, the *Per2:luc* reporter activity under a temperature cycle (**Figure 2d**) was very similar to that seen in mouse skin fibroblasts, but with a remarkable 9-12h phase jump when cells were released back into constant conditions. This surprising behaviour was paralleled by unusual patterns of *Bmal1:luc* reporter activity (**Figure 2e**). Here, unlike the case in mouse fibroblasts, we observed an immediate alignment to the temperature cycle (**Figure 2f and Suppl. Fig. S2b**), with suppression by rising temperatures similar to the *Per2:luc* construct. When these cultures were released back into constant temperature conditions *Bmal1:luc* and *Per2:luc* returned to an approximately antiphasic relationship with one another within the first cycle of free-running. Unlike the case for mouse fibroblasts, cultures that start the experiment in antiphase to one another still had quite distinctive phases (Bmal1:*luc* Ψ = 3h 38min ± 20min; t-test, p = 0.0004, t = 11.18, df = 4; *Per2:luc* Ψ = 2h 24min ± 36min; t-test, p = 0.0168, t = 3.951, df = 4).

We wondered if the observed acute sensitivity of *Bmal1:luc* to temperature was an artefact of the reporter construct. To test this, we cultured untransduced mouse and reindeer skin fibroblasts in six-well plates before synchronising them with DEX and making serial collections over the first temperature cycle at a 4h sampling resolution. We then measured the abundance of mouse and reindeer *Per2* and *Bmal1* mRNA during the time course using qPCR. In mouse skin fibroblasts mouse *Per2* and *Bmal1* mRNA levels peaked in distinct phases during the temperature cycle (**Figure 2g**), whereas reindeer *Per2* and *Bmal1* mRNA levels were overlapping in phase (**Figure 2h**). These data are consistent with the reporter activity and provide independent evidence that, in contrast with mouse skin fibroblasts, temperature has a strong and immediate impact on both *Bmal1* and *Per2* gene expression in reindeer skin fibroblasts.

### Mutation of RORE elements in a *Bmal1* promoter reporter produces reindeer-like reporter rhythms in a mouse cell context

The REV-ERB and ROR families of nuclear hormone receptors are principal regulators of circadian *Bmal1* expression. These factors activate (RORs) or repress (REV-ERBs) transcription through two conserved RORE sites located in the *Bmal1* proximal promoter (Abe et al., 2022; Akashi and Takumi, 2005; Guillaumond et al., 2005; Preitner et al., 2002; Ueda et al., 2002), and, in the mouse, this stabilises circadian rhythm expression (Abe et al, 2022).

We showed that *Bmal1:luc* in reindeer fibroblasts and reindeer *Bmal1* mRNA abundance is highly temperature-sensitive (**Figure 2**). The proximal promoter of reindeer *Bmal1* contains both ROREs deemed necessary for circadian transcription (**Suppl. Fig. S4**), and reindeer RORS and REV-ERBs show strong sequence homology across the functional domains required for their transcriptional activator/ repressor activity (**Suppl. Fig. S5)**. These findings may suggest that the effect of temperature on *Bmal1* promoter activity is independent of the effect mediated via the circadian clock and its known early temperature-responding elements such as *Per2* (Buhr et al., 2010; Saini et al., 2012; Tamaru et al., 2011).

To test this hypothesis, we mutated both RORE elements in the *Bmal1:luc* reporter (creating a new reporter: *Bmal1-mutated:luc*) (**Figure 3a**). To confirm that the circadian clock machinery was decoupled from our new *Bmal1-mutated:luc* reporter we compared the reporter activity of the *Bmal1:luc* and *Bmal1-mutated:luc* reporters following DEX synchronization at a constant ambient temperature. As expected, transduction of U2OS cells (a human-derived stable cell line and common *in vitro* molecular clock model), mouse skin fibroblasts and reindeer skin fibroblasts with the *Bmal1-mutated:luc* reporter showed a profound loss of rhythmic reporter activity compared to cells transduced with the wild type *Bmal1* reporter (**Suppl. Fig. S3 a-c**). We next ran another temperature entrainment experiment but this time comparing the *Bmal1:luc* reporter to the *Bmal1-mutated:luc* reporter in mouse, reindeer and U2OS cells. The responses of the wild-type *Bmal1:luc* reporter in mouse and reindeer skin fibroblasts (**Figure 3b**) were consistent with our previous experiment (**Figure 2**). Interestingly, whereas the *Bmal1-mutated:luc* reporter showed no cyclicity in either mouse, reindeer or U2OS cells following DEX synchronisation, it showed a strong thermal response with rising temperatures causing a dramatic suppression of bioluminescence and falling temperatures increasing bioluminescence (**Figure 3b and Suppl. Fig. S3d**). Moreover, the relative phases of the bioluminescence profiles relative to the temperature cycle for the *Bmal1-mutated:luc* reporter were strikingly similar to those seen for the wild-type *Bmal1:luc* reporter in reindeer fibroblasts (**Figure 3c**).

**Figure 3.**
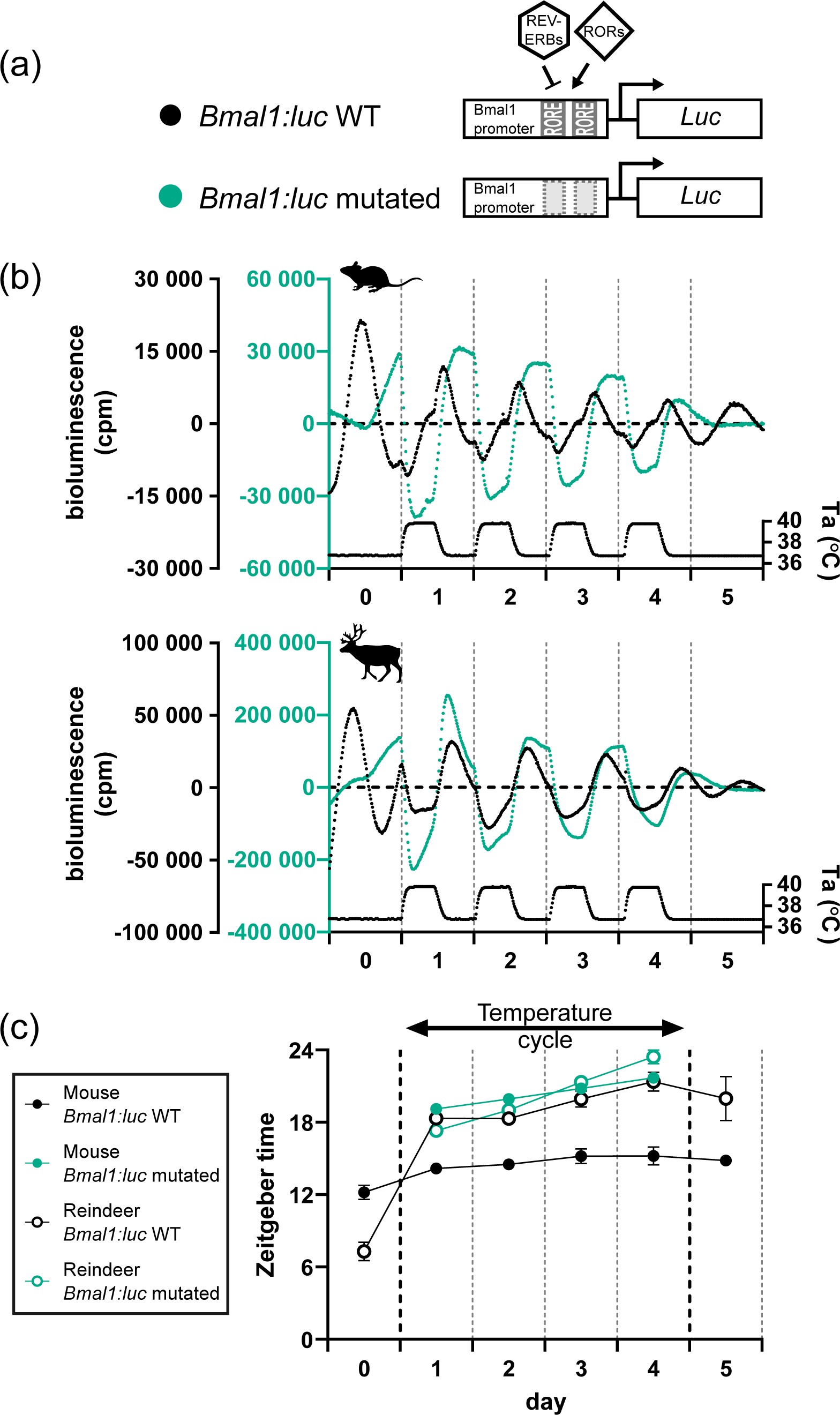
RORE mutation of the *Bmal1:luc* reporter and responses under temperature cycles. (a) RORE sites in the *Bmal1* promoter facilitate the response of REV-VERBs and RORs on *Bmal1* expression. Both RORE sites in the promoter reporter sequence were rendered non-functional by mutation (WT: un-mutated). (b) The mutated and un-mutated reporters were transduced into mouse and reindeer fibroblasts and were measured under an ambient temperature (Ta) cycle for four days. (c) Peak expression was determined by the CircWave programme and plotted for both species and both promoter reporters.

## DISCUSSION

Taken together our data shows that reindeer skin fibroblasts have a circadian rhythm that can be synchronised with DEX, is temperature-compensated and undergoes partial entrainment following 24h temperature cycles. We show that reindeer fibroblasts have a comparatively short free-running period compared to mouse cells and that *Bmal1* has heightened sensitivity to temperature in reindeer compared to mouse fibroblasts. We argue that this might be due to weak coupling between the core and secondary loop in the reindeer molecular clock.

Previous work by Lu et al (2010) reported that the reindeer molecular clock was non-functional. The striking differences between our results and those presented by Lu et al. (2010) were unexpected, especially as we ensured minimal differences between our experimental approaches. The reindeer skin biopsies were both collected from animals that originated from a herd held at the University of Tromsø, Norway (69°N) and the culture conditions and synchronisation protocols between the studies were identical. The *Per2:luc* and *Bmal1:luc* reporters used in our study are of different origins, but ultimately, both sets of promoter reporters contain the key regulatory elements that allow mouse skin fibroblasts to drive rhythmic, phase-specific transcription. In contrast to the original study, we included an additional blasticidin selection step to ensure that all fibroblasts expressed the reporter constructs, However, it is unlikely that the previously reported arrhythmicity of reindeer fibroblasts is caused by a lower lentiviral transduction efficiency. Under such a scenario, weak rhythmicity would also be expected in the positive control, i.e. mouse fibroblasts. It is possible that variable cell density may explain the differences between our studies. We ensured that our cells were confluent before synchronisation as low cell density have been reported to affect circadian rhythmicity of skin fibroblast cultures (Noguchi et al., 2013). Although the use of confluent cultures in PMT work is a standard procedure, the confluency of the reindeer skin fibroblasts used in the Lu *et al* (2010) paper was not stated.

Although reindeer skin fibroblasts satisfy key criteria of circadian rhythms, we identify several distinctions between our mouse and reindeer model systems. Most notable were the data collected during our daily temperature cycle experiments. Here, the expressions of both *Bmal1* and *Per2* were temperature-sensitive in reindeer skin fibroblasts, leading to an unusual in-phase expression profile between the genes. We also observed a low amplitude and comparatively weak circadian alignment of the reindeer cultures once they were released back to constant temperature. This likely reflects the re-establishment of the cell-autonomous circadian oscillators within the reindeer skin fibroblast population. Here, the interaction of the positive (*Bmal1*) and negative (*Per2*) regulators of the core molecular clock resolve from their atypical in-phase state to their typical anti-phase relationship evident as a large phase shift in *Per2*. The dampened amplitude might reflect phase variation between individual cells in the newly established circadian phase measured at the level of the whole cell population in a dish (Welsh et al., 2004).

Intrigued by the unusual regulation of *Bmal1* by daily temperature cycles in reindeer skin fibroblasts, we turned our attention to the molecular processes that underpinned the striking differences in daily temperature cycles between mouse and reindeer skin fibroblasts. To investigate the role that RORE-mediated transcription plays in temperature resetting, we experimentally uncoupled the circadian clock machinery from the *Bmal1:luc* reporter by mutating the two canonical RORE elements. Our experiments using this construct revealed a strikingly similar response to temperature cycles in mouse and U2OS cells as in reindeer with the wild-type reporter, namely an immediate phase shift with a trough at temperature rise (**Figure 3**).

These data show that RORE mutation, and therefore removal of the ROR / REV-ERB interaction at the *Bmal1* promoter, leads to clock-independent temperature sensitivity of the reporter. It is clear, however, that endogenous circadian ROR / REV-ERB expression does regulate the *Bmal1:luc* reporter in reindeer skin fibroblasts as there is a robust circadian rhythm under constant conditions in reindeer skin fibroblasts which is disrupted by RORE mutation.

This leads us to suggest a model in which a species difference in the relative weighting between direct, RORE-independent effects and indirect RORE-mediated, circadian effects of temperature accounts for the different responses seen in temperature entrainment experiments (**Figure 4**). Under this model, in mouse fibroblasts, the ROR / REV-ERB transcription factors strongly couple *Bmal1* expression to the circadian clock, and it is this, probably via direct effects of temperature on *Per2* expression (Buhr et al., 2010; Tamaru et al., 2011) which mediates the entrainment response to temperature cycles. In contrast, in reindeer fibroblasts, circadian clock-mediated effects of temperature on the ROR / REV-ERB transcription factors are seen as exerting only a weak regulatory effect on *Bmal1* transcription, while direct, RORE-independent effects of temperature dominate; this effectively weakens the coupling between the primary and secondary loops of the reindeer clockwork. Interestingly, mathematical modelling of the circadian clock predicts that a weakened influence of the secondary loop on the core loop leads to shortened free-running periods (Yan et al., 2014), a prediction that is consistent with the comparatively short free-running period of the reindeer compared to mouse skin fibroblasts.

**Figure 4.**
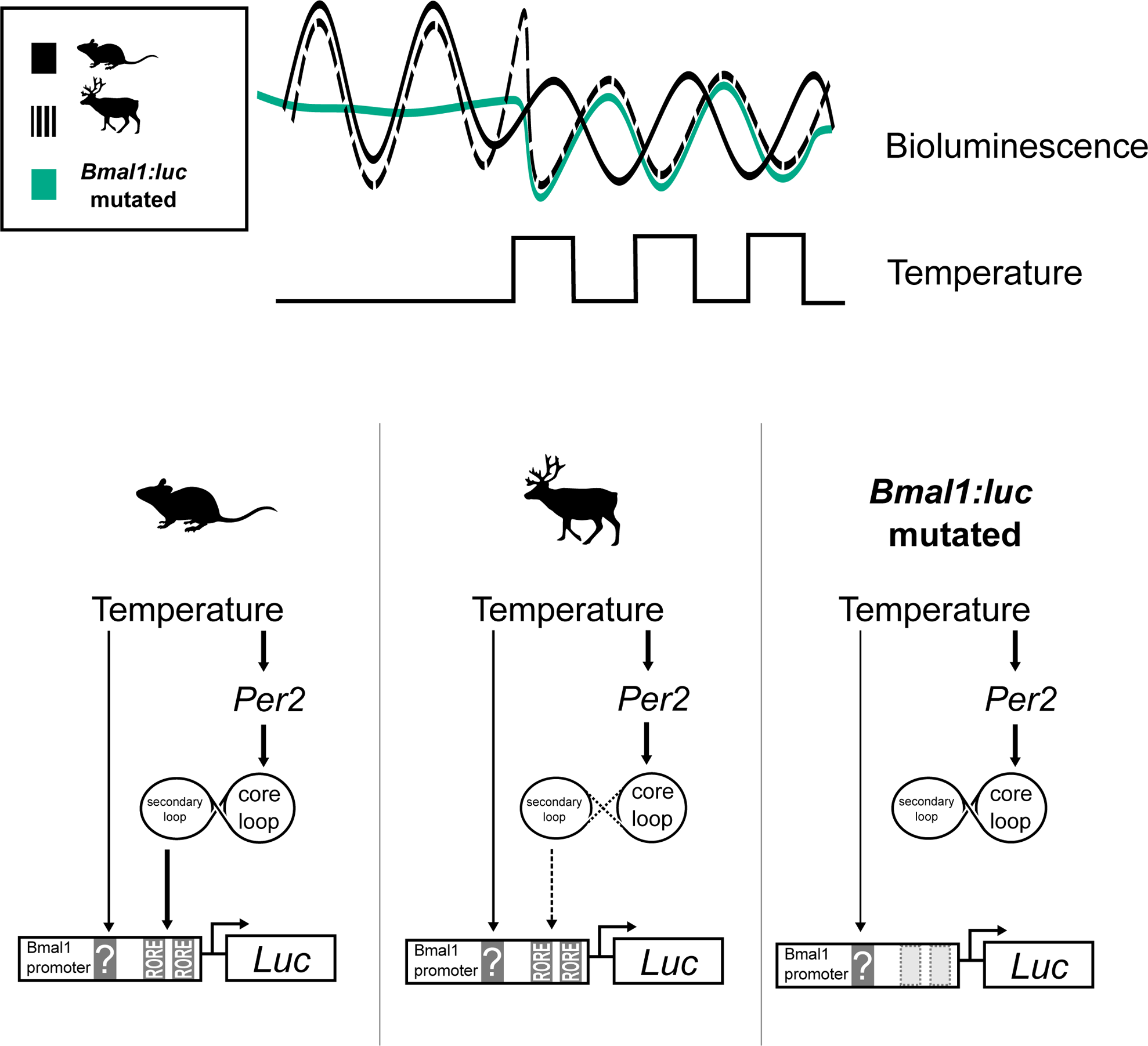
In mouse fibroblasts, the *Bmal1:luc* reporter entrains gradually to an ambient temperature cycle eventually resuming its approximate antiphase relationship with *Per2* expression, suggesting that the *Bmal1*-specific temperature effect is dominantly mediated via the circadian clock. In reindeer fibroblasts, *Bmal1* is acutely responding to temperature cycles and moves into phase with *Per2* expression. We suggest that a weak coupling between circadian loops reduces the effect of temperature via the RORE elements on the *Bmal1* promoter, instead an unknown RORE-independent pathway drives *Bmal1* expression during temperature cycles in reindeer fibroblasts. Supporting this model: when we experimentally decoupled the circadian clock from the *Bmal1:luc* reporter by mutations of the ROREs we rendered the promoter-reporter acutely temperature-sensitive which led to a reindeer-like phase response under temperature cycles.

Our experiments leave open the question of how coupling between the reindeer core and secondary loops has become weaker than is the case in mice. We could find no obvious differences in either proximal *Bmal1* promoter organisation or in the major DNA binding, ligand binding and protein-protein interaction domains in the REV-ERBs and RORs. It is possible that a recently described a reindeer-specific mutation in the PER2 protein, which affects its interaction with CRY1 (Lin et al., 2019), is involved in the phenomena we describe, and genetic manipulation of this site is a desirable future objective.

The whole-organism implications of a reindeer-type compared to a mouse-type circadian oscillator are difficult to predict. However, given that thermoenergetic demands impact the circadian organisation of 20 g nocturnal rodents radically differently compared to 100 kg diurnal ungulates of the Arctic (Hazlerigg and Tyler, 2019; van der Vinne et al., 2014), we speculate that our results reflect unappreciated diversity in the coupling of heat-sensitive signal transduction to circadian entrainment and metabolism.

## Acknowledgements

The authors would like to thank our animal technicians Hans Lian, Hans-Arne Solvang and Renate Thorvaldsen. Only their dedication and hard work allows us studies on our fascinating Arctic model organisms. We also thank Jaione Simón-Santamaría and Karolina Szafranska for the mouse skin biopsies and Barbara Tomotani for useful discussions on the manuscript. We also thank Fredrik Markussen for his circular-plotting skills.

## Data availability statement

pLV6-mPer2-luc generated in this study have been deposited to Addgene, Plasmid #212035. PMT and qPCR data are deposited in DataverseNO: https://doi.org/10.18710/YCURBQ

