## Supplementary Figure Legends for "The reindeer circadian clock is rhythmic and temperature-compensated but shows evidence of weak coupling between the secondary and core molecular clock loops"

**Table S1.** Sequences for all primers used. Underlined sequences refer to mutation sequences. Mutation was done with the Site-directed mutagenesis (SDM) kit.

| | Primer | Primer sequence (5'-3') |
|---|---|---|
| SDM mutation primers: Inserting BamHI cutting site | forward | CGACGGTATCGGATCCCGAGCTCAAG<br>CTTCG |
| | reverse | CGACGGTATCGGATCCCGAGCTCAAG<br>CTTCG |
| SDM mutation primers: RORE mutation | RORE1 forward | GGGGAGCGGATTGGTCGGAAAGTGT<br>ACGTG<br>TGGTGCGACATTTAGGGAAGG |
| | RORE1 reverse | CCTTCCCTAAATGTCGCACCACACGT<br>ACACTT TCCGACCAATCCGCTCCCC |
| | RORE2 forward | GACATTTAGGGAAGGCAGAAAGTGTA<br>CGTGG<br>GACGGAGGTGCCTGTTTACCC |
| | RORE2 reverse | GGGTAAACAGGCACCTCCGTCCCACG<br>TACAC TTTCTGCCTT CCCTAAATGTC |
| qPCR primers | Mouse Ppib forward | AAGTCACAGTCAAGGTATAC |
| | Mouse Ppib reverse | TAGCCAAATCCTTTCTCTC |
| | Mouse Bmal1 forward | CTGAAACACCTAATTCTCAG |
| | Mouse Bmal1 reverse | CATTCTGGCTATAATTGAGG |
| | Mouse Per2 forward | CACAAAGAACTGATAAGGAC |
| | Mouse Per2 reverse | CTGGTAGTACTCCTCATTAG |
| | Reindeer Ppib forward | CTGAGAATTGGAGATGAAGA |
| | Reindeer Ppib reverse | GGAATTTGCTGTCTTTGTAG |
| | Reindeer Bmal1 forward | TGAGTATTTCCATCAAGACG |
| | Reindeer Bmal1 reverse | ATGAAACTGAACCACCGA |
| | Reindeer Per2 forward | GCCATCATTATCTGCAAG |
| | Reindeer Per2 reverse | TCCAGAGGTATTTCTTAGTC |

**Figure S1.** All replicates of mouse and reindeer fibroblast transduced with *Bmal1:luc* under different constant ambient temperatures (n=6 cultures for each species).

**Figure S2.** (a) All replicates of mouse and reindeer fibroblast transduced with either *Bmal1:luc* or *Per2:luc* and measured under an ambient temperature cycle. Furthermore, cultures for each respective gene and species were DEX-synchronized 12h apart from each other, effectively resulting in eight experimental groups: Mouse Bmal1 + 0, Mouse Bmal1 + 12, Mouse Per2 + 0, Mouse Per2 + 12, Reindeer Bmal1 + 0, Reindeer Bmal1, Reindeer Per2 + 0 and Reindeer Per2 + 12 (n = 3 for each group). Zoomed-in versions of the baseline-corrected bioluminescence data are provided for *Bmal1:luc* and *Per2:luc* in reindeer fibroblasts after the temperature cycle. (b) Data of peak expression (centre of gravity) for each gene, species and day of the temperature cycle experiments. Data is plotted in circular plots to visualize the phase relationships. Data corresponds to **Figures 2c and 2f**.

**Figure S3.** Three different cell lines (U2OS, mouse fibroblast and reindeer fibroblasts) were either transduced with the *Bmal1:luc* or the RORE-mutated *Bmal1:luc* reporter. (a) Figures in the first row show the raw data plotted on a common axis to visualize absolute *luciferase* expression levels between the reporters. (b) Figures in the second row show the same data plotted on different axes to visualize rhythmicity or the lack thereof. (c) Figures in the third row show baseline corrected data. (d) Figures in the fourth row show all replicates of wildtype and the mutated promoter reporter under the temperature cycle (n = 4 cultures for each respective reporter and cell line).

**Figure S4.** Sequence alignment between the mural *Bmal1* promoter of the *Bmal1:luc* reporter (pLV6-Bmal1-luc) and the corresponding reindeer sequence.

**Figure S5.** Amino acid sequence alignment of RORs and REV-ERBs between mouse and reindeer.
