## Supplementary figures and images for "The reindeer circadian clock is rhythmic and temperature-compensated but shows evidence of weak coupling between the secondary and core molecular clock loops"

### Supplementary Figure 1

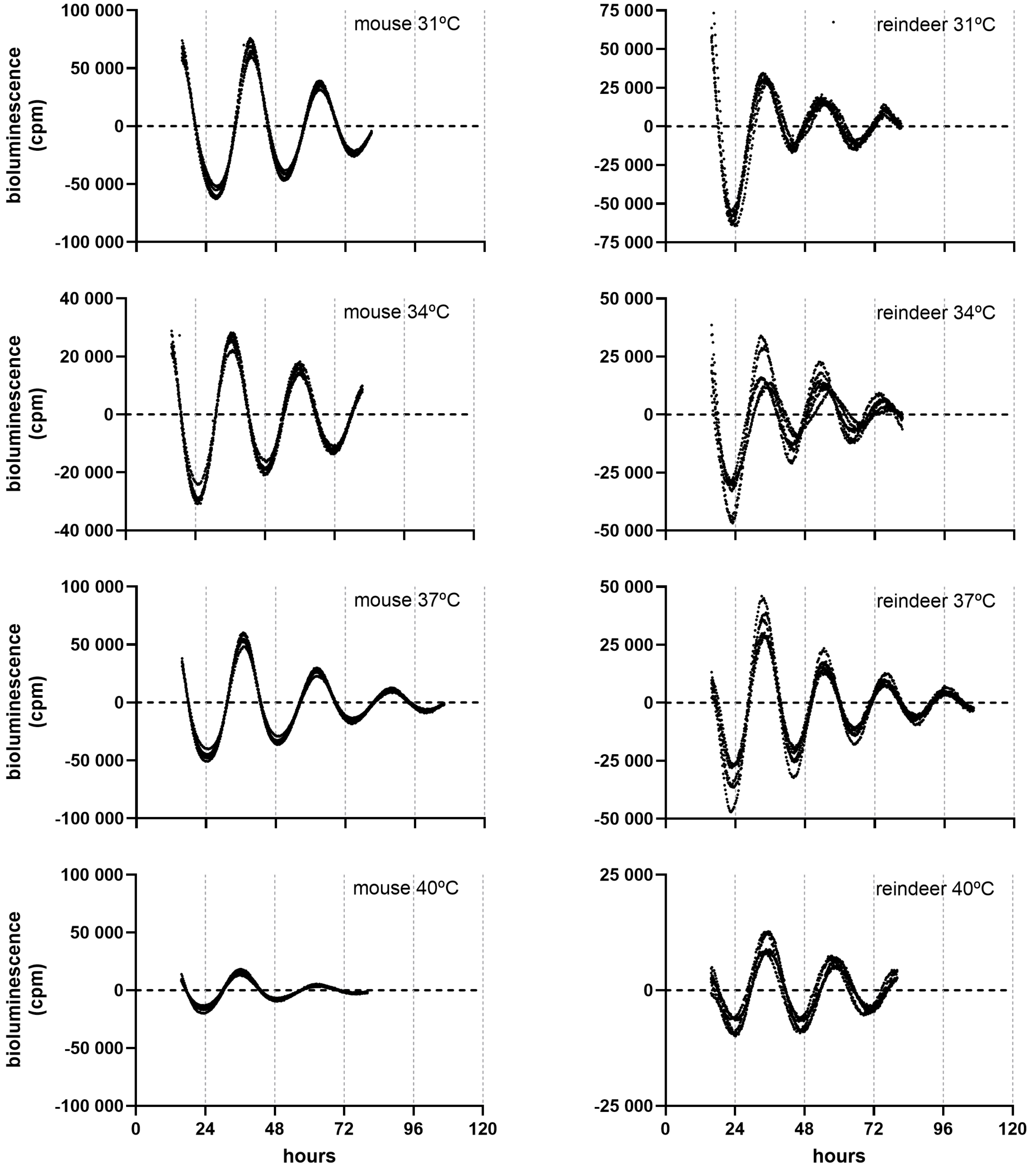

### Supplementary Figure 2

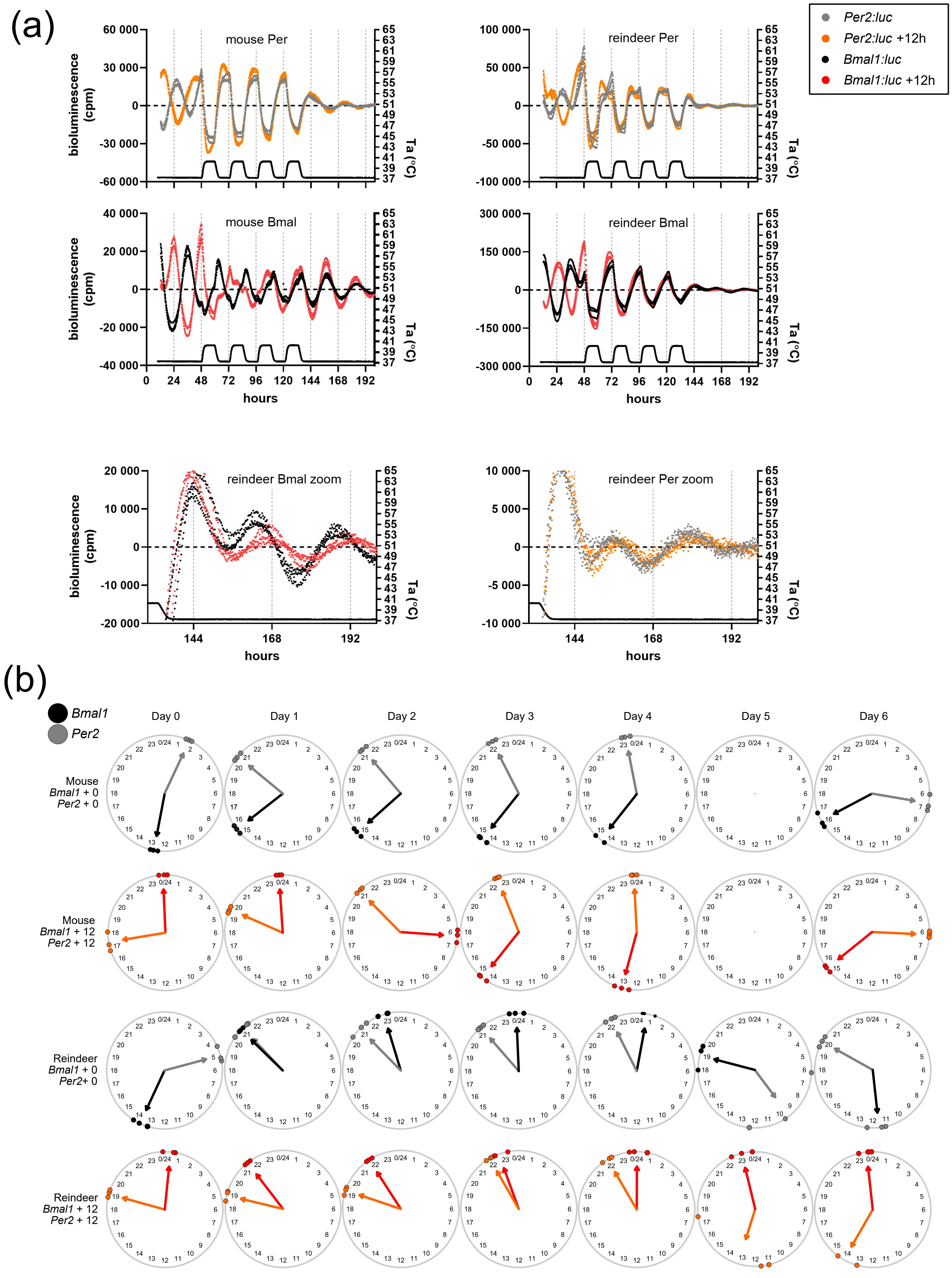

### Supplementary Figure 3

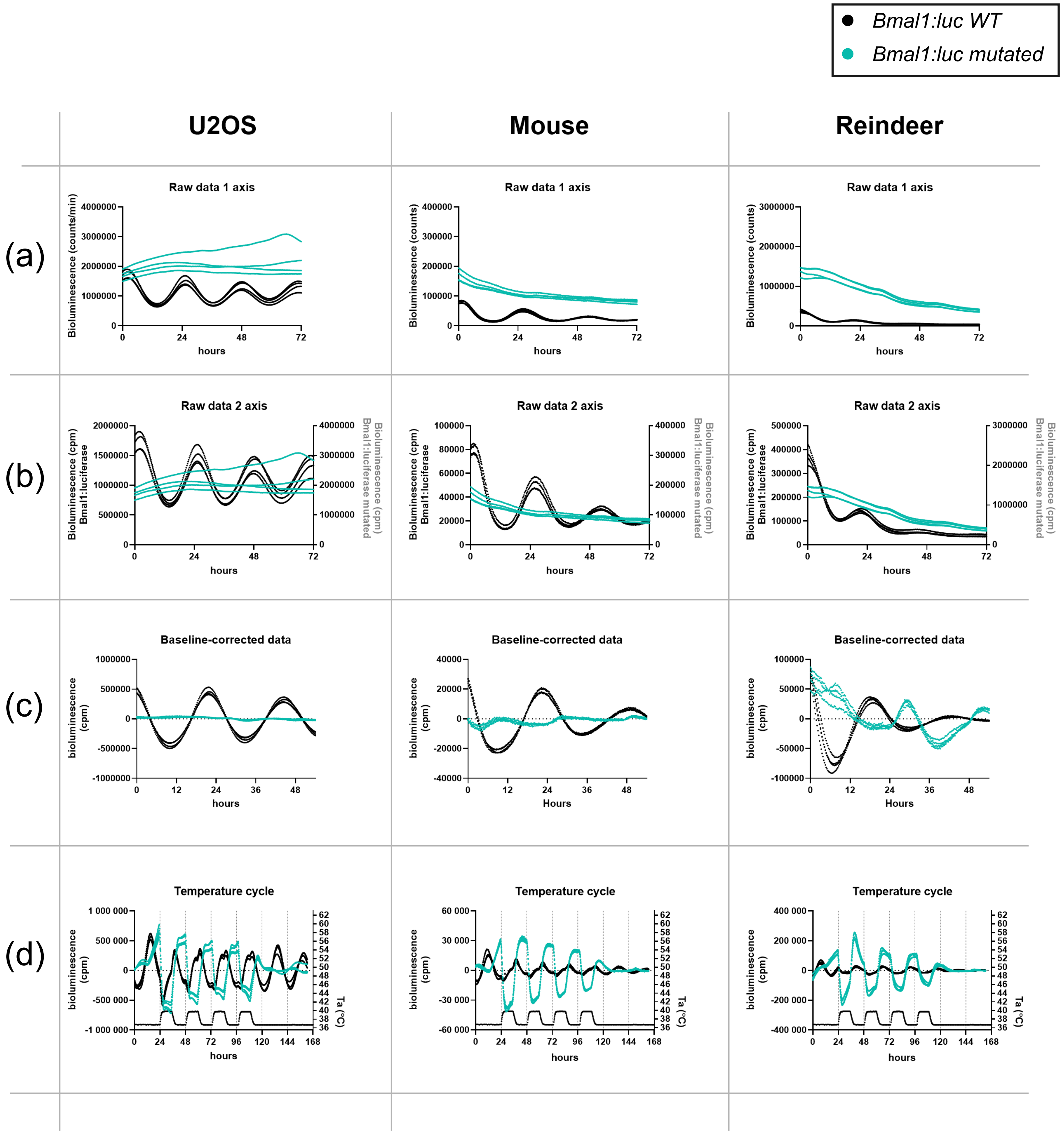

### Supplementary Figure 5

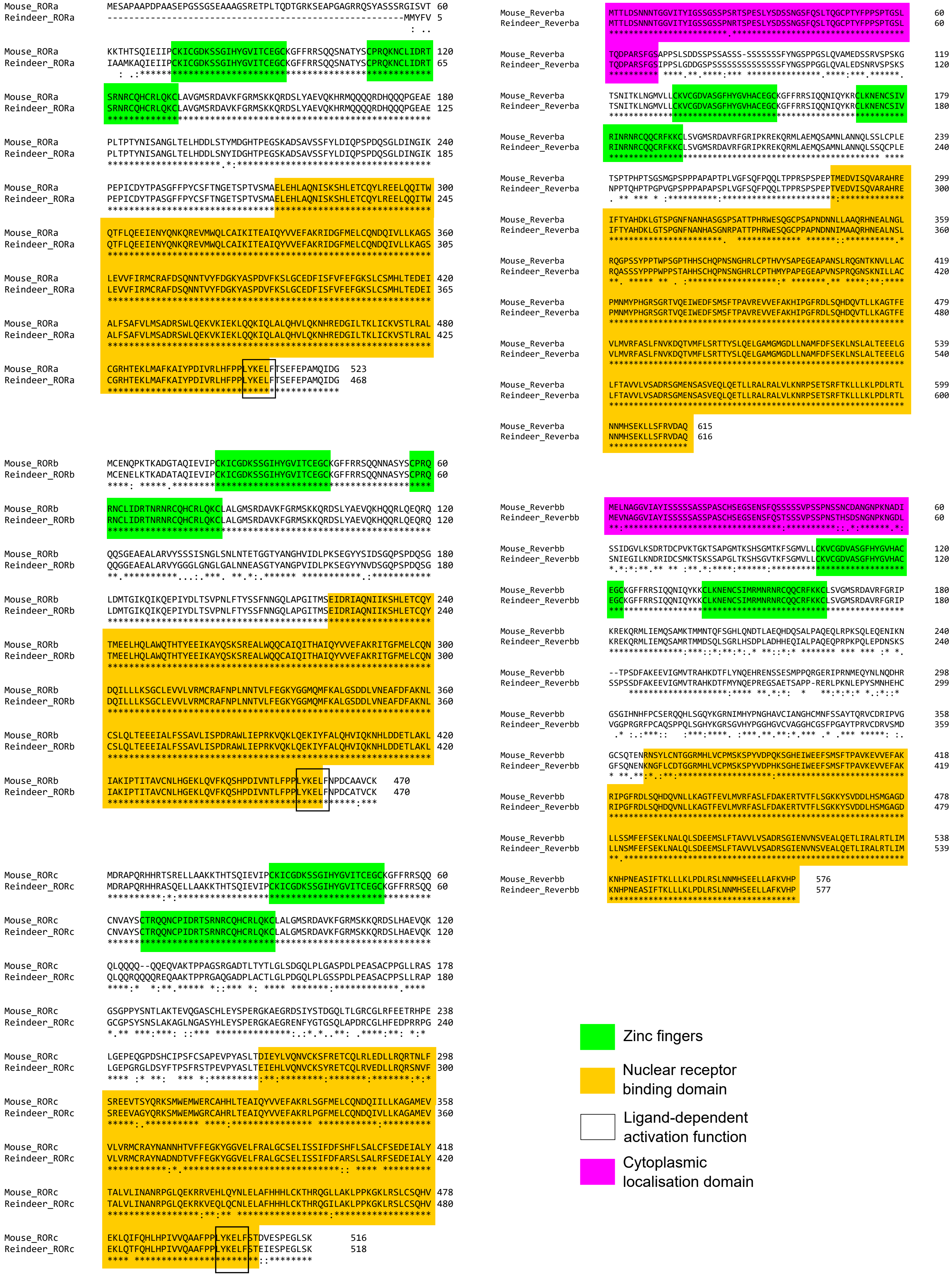
