## Supplementary Figure 4 for "The reindeer circadian clock is rhythmic and temperature-compensated but shows evidence of weak coupling between the secondary and core molecular clock loops"

5' upstream sequence

```
reporter      GGGCTACAACAGAACAACTAACCACCCAAACAAACAAC      38
reindeer      GTGTTAAAGGCCCTCTGTTAAG-GGCTTGAGATGCAAA      59
              * * ** *      *   ***    *   * *  ***

reporter      TTTGCAACTTTTCTTTCTTTCCCG-CTCC--TTTGA--TACTTTTGCAAAGTACATTTGC      93
reindeer      AATAAGATTCTTCGTCCTCTCAGTACTCAAGTTGTATCTTGTTACGAAACGTTTATTTTC      119
                *   * * *** * ** **    ***   **  *  *  **  * ** **  **** *

reporter      TAGGGCCATGTCCATAACATGTAATAGAATCTTGCTCAGGTCTCACCCTGTGGCCCCGGC      153
reindeer      TGCAACTATTTGTTTCCTTCCGTCTCAGTCCTTGGACAGCTTTTCTACGGTAGAAACAGC      179
              *    * ** *   *         *     ****  *** * *    * ** *   * **

reporter      TAGCTCAGTACTCGCGATTATGCCCCTGCCTCAGCTTCTTGAG--GGTTGGAATTACAGA      211
reindeer      ---------CACTGGGAAGTT--GTCTGCCACACAGTGGTGCGGCCACTGAAAGGAAAAG      228
                           * **   *    ***** **   *  ** *     ** **  * *

reporter      C---TACGCCACCACCCCCGGGAGACTTCTCTTCCGATCGGGAATGACAGCTCCACTGGA      268
reindeer      CACTGGAATCGCTCCAGGCTTGAGTTTTCTCTTTTGAACAAGAAAGACAGCTCCAATCGG      288
              *        * *  *   *  ***  *******  ** *  *** ********** * *

reporter      -CGCTTGGAGGTCAAGAGAAAAAAAAAAAAAAAAAAAAAAAAGAACGCGCGATCCGGTTA      327
reindeer      ATGCTTGGAGGTCAGGACGAACGAAAAGAACGTAAGAA----------------------      326
                ************ **  **  **** **   ** **

reporter      GTCGTCGGAAGGAAAAGTCTTCTAAATGCGCTGGCTATTAGCGCTGTGGTTCCGACTAGG      387
reindeer      -CACTTGAATGGAAAAACCTTCTAGAAATACCAAGTATTTCTTCTTCGTTTGGACTTTGG      385
                  * * * ******  ****** *    *    ****    **  * **     * **

reporter      ACAAGCCAGGGGTTTCAGGAAGGTTCTGTCACTCTGTGTTCCTAATATGTGGTTTCCTAG      447
reindeer      CAAACCAGGATATTTTGGGAAGGGTCTGACACTATGTT--CCT-AAAACTGGTTTCCTAG      442
                ** *  *   ***  ****** **** **** ***   *** * *  ***********

reporter      ATGGAGCTGGAGAGGGATGGGCGAAGAGATGCGGCGTTTTCTCAGTGTTCTGCGAGCCAG      507
reindeer      ATGGAGTTGGAGGTAGGGGAACGAGGAAATGTTGC-TTTAGTCAGGATGCCCCGAGTGAA      501
              ****** *****   *  *  *** ** ***  ** ***  ****  * *  ****  *

reporter      GAAGGGGACGGAGGAGGAGGGAGAGCTGAGAGGCAGTCACTAGCCAGGGACGCGGCTGAG      567
reindeer      AAAGGGCAAGGAAGAGGAGTGAGATACGTGACCCAGAAGCAAGTCAGGAACCTAGAGAAG      561
               ***** * *** ****** ****   * **  ***   * ** **** **   *   **

reporter      GAGCAGGGGACAACGGCG-----AGCTCGCAGAGTC-CGCAACGCAGTGGCCTCAGCGAG      621
reindeer      AAGGACATCGCAGGCGACTGCAAAGAGTGCGGCGTCTGGACCCGCAGTGGCCAAGGCTAA      621
               ** *     **   *       **   ** * ***  *   **********   ** *

reporter      CTTTAGACCTGAGGG-GGAAAACCGAGGGCTGGGGCAAGAAATCCACAGAGCGTGCCAAT      680
reindeer      CGGGAGAGCAGGGGAAAGAGAGAGAACAGAGGCGAGGAAATCCAGGGAAAACGCGTCGAT      681
              *   *** * * **   ** *    *  *  * *   * *       * * ** * * **

reporter      TGGTCCACTCCTCGGGGCGTGTGCTTCTGTGCGCCAAATGATTGGTGAAGGGAAAGTAGC      740
reindeer      TGGTCCTCTCCTTGGGGCGTGTGCTTCTGTGCGCCAAACGATTGGTGGCGGGAAAGTAGC      741
              ****** ***** ************************* ********  ***********

reporter      AGGTAAACCAGCCCTGCCGT-CTTGCCATTGGTCAGAGGCTTTCCTATCGGTCACTCGGT      799
reindeer      AGGTAAACCGGCCCTCGCCGCCTCGCCATTGGTCAGCGGCCTCTCTTGATGCCACACAAC      801
              ********* *****  *   ** ************ *** *  **    * *** *

reporter      TGGCTAGCCTAACGCAGAGCAGAACGCGAATTGGTTTGGGTTGTCC-GCCAAGACAACTC      858
reindeer      TGGCTATCAGGCCGCCGAGCCGCGCGCGGATTGGCCGGGGGCGGCCGCCGAGACTATCTC      861
              ****** *    *** **** *  **** *****   ***  * **  * *    * ***

reporter      CGTTCGCTCTCTCTGATTGGCTAACGGGAAGAGGCAGGTATCCGGGCTGCGCGGCTCCTC      918
reindeer      TCTTCGGGCTCTAGGATTGGCTGGAGGGAAGAGGCAGGTATCCGGGCTCCGCGACTCCTC      921
                ****  ****  ********   *********************** **** ******

reporter      CATTGGTGGGCGGGGAAGGGGGGTTGGGCACAGCGATTGGTGGGCGGGGGGCCGGGCCTG      978
reindeer      CATTGGTGGGCGGGGAAGGGGGGTTGGGCACAGCGATTGGTGGGCGGGGGGCCGGGCCTG      981
              ************************************************************

reporter      GGCCGGCGGGGAGCGGATTGGTCGGAAAGTAGGTTAGTGGTGCGACATTTAGGGAAGGCA      1038
reindeer      GGCCGGCGGGGAGCGGATTGGTCGGAAAGTAGGTTAGTGGTGCGACATTTAGGGAAGGCA      1041
              ************************************************************

reporter      GAAAGTAGGTCAGGGACGGAGGTGCCTGTTTACCCGCGCCGGACTTGCCGCCGCCGC      1098
reindeer      GAAAGTAGGTCAGGGACGGAGGTGCCTGTTTACCCGCGCCGGACTCACCGCCGCCGC      1101
              *********************************************  *************

reporter      CGCGGGATC 1107
reindeer      CGCGGGATC 1110
              *********
```

3' downstream sequence

RORE 1

RORE 2

Figure S4
